# Adera: A drug repurposing workflow for neuro-immunological investigations using neural networks

**DOI:** 10.1101/2022.07.14.500072

**Authors:** Marzena lazarczyk, Kamila Duda, Michel-Edwar Mickael, Agnieszka Kowalczyk, Mariusz Sacharczuk

## Abstract

Drug repurposing in the context of neuro-immunological (NI) investigations is still in its primary stages. Drug repurposing is an important method that bypasses lengthy drug discovery procedures and rather focuses on discovering new usage for known medications. Neuro-immunological diseases such as Alzheimer’s, Parkinson, multiple sclerosis and depression include various pathologies that resulted from the interaction between the central nervous system and the immune system. However, repurposing of medications is hindered by the vast amount of information that needs mining. To challenge the need for repurposing known medications for neuro-immunological diseases, we built a deep neural network named Adera to perform drug repurposing. The model uses two deep learning networks. The first network is an encoder and its main task is to embed text into matrices. The second network we explored the usage of two different loss function, binary cross entropy and means square error (MSE). Furthermore, we investigated the effect of ten different network architecture with each loss function. Our results show that for the binary cross entropy loss function, the best architecture consists of a two layers of convolution neural network and it achieves a loss of less than 0.001. In the case of MSE loss function a shallow network using aRelu activation achieved an accuracy of over 98 % and loss of 0.001. Additionally, Adera was able to predict various drug repurposing targets in agreement with DRUG Repurposing Hub. These results establish the ability of Adera to repurpose with high accuracy drug candidates that can shorten the development of the drug cycle. The software could be downloaded from https://github.com/michel-phylo/ADERA1.

## 1. Introduction

Drug repurposing represents a lifeline for the immunology drug industry. In recent years, the rate of novel drug development has been rapidly declining^1^. This phenomenon was accompanied by a sharp rise in drug development costs^1^. Currently, it could cost about one billion dollars to produce one single novel drug ^2^. Drug repurposing constitutes a viable alternative to conventional drug development techniques. Repurposing drugs that have been already legalized for human usage can substantially reduce the costs accompanying the primary stages of drug discovery. Moreover, repurposing novel drugs eliminates the delay faced by de novo drug development ^3^. At present numerous repurposed medications are extensively used such as Azathioprine which was commonly used for rheumatoid arthritis, and currently, it is used for renal transplant^4^. Digoxan was originally given to patients who suffer from cognitive heart disease and currently, it is used to treat cancer, similarly, arsenic was known as a drug for syphilis and now it is being used to treat leukemia. We previously repurposed zileuton which was originally used as an inhibitor of the 5-lipoxygenase as an Nrf2 activator to treat depression ^3^. By using drug repurposing strategies the total time needed for drug development can be shortened significantly.

Drug repurposing for immunology disorders using text mining is still far from perfection. Currently identifying drug targets could be done through three main approaches, machine learning, network-based, and text mining ^5^. Machine learning approaches are based on grouping of compounds and diseases. However, they suffer from inherent bias, typically because of the production of negative samples^6^. Although network approaches are widely used to predict drug-disease interaction, they are unaware of the molecular relationship pertaining to the investigated diseases. Text mining approaches have proven to be a reliable alternative for other drug repurposing methods. However, current text mining methods are still constrained by the limitations of natural language processing tools^6^. Deep learning for text mining approaches has overcome the traditional limitations of text mining approaches. However current deep learning approaches are unaware of the relevance or the context of the text^3^.

In the current study, we used two different neural networks in a single workflow to increase the relevance of compounds repurposed as well as decrease false positives. The model accepts a pathway/disease and then scans Pubmed for relevant documents. Each relevant document is embedded into a matrix using an encoder. After that, each embedded matrix is processed using a “relevance” neural network that predicts the relationship between each row (i.e., a sentence) in the matrix and the pathway/disease in question. Subsequently, the sentences containing drugs are sorted based on their relevance. We crossed validated ADERA1.1 performance against a gold standard dataset (Drug Repurposing Hub, Broad Institute dataset)^7^. In silico validation using a case study was done as follows. We investigated repurposing anti-oxidants compounds that could be used to regulate Th17 cells in depression. Previously, we have found that Th17 infiltrates the blood-brain barrier through a paracellular route causing depression-like behavior^8,9^. The main objective of the *in-silico* validation step was to find anti-oxidant drugs that could be used to reduce the effect of Th17 in depression. In order to be counted as a hit the compound has to be an antioxidant, has an inhibitory effect on Th17, is known to treat depression, and is capable of passing the BBB. Finally it had to score at least 4 out of 5 on Lipinski’s score. Out of the top ten compounds generated by our software two were shown to be true positives.

## 2. Methods

### 2.1 Overview of the workflow

The workflow of the program is as follows. The first phase of the program covers the aim of downloading pdfs for processing in downstream steps. This phase consists of three further steps. The first step objective is to fetch the Pubmed IDs related to the search query. This is accomplished by using the Pubmed fetcher function available through the Metapub python library. This step takes in the first input query (e.g., what are the currently used colorectal cancer drugs (CRC)) and searches for the recent PubMed articles that discuss them. The number of articles returned to the user can be adjusted through the options menu in the name window. After that, in the second step, the program fetches the abstracts of the retrieved PubMed Ids. The third step of this phase is the extraction of the keywords from the abstracts and it is achieved through the usage of the python library Keybert. Following that the user has an option to choose which pdfs to process with or proceed automatically. The fifth step in this phase is the downloading and cleaning of the pdfs and this is done using the fetch pdfs library.

The second phase of the software is the parsing of each pdf into separate sentences and storing all the pdfs in a single database using the JSON format. This phase uses several python libraries which are essential for text extraction including Tika, ntk, and JSON. The third phase of the software compromises two neural networks. The first one is an autoencoder and it is known as Google universal encoder text^10^. This network’s main function is embedding each sentence of the extracted sentence from each pdf into a 1× 512 matrix. The second step in this phase is the relevance network which is achieved using various Keras architectures in sequential form. We constructed two different networks Adera1 and it used an Adam optimizer and loss of function of binary_crossentropy. The second one is Adera1 and it used an Adam optimizer and loss of function MSE (regression). We tried ten different network layers for both networks.

### 2.2 Performance metrics

To measure the performance of the second neural network model, we trained it on the database SciQ Dataset, which consists of three sub-datasets (training, validation, and testing). The Dataset was downloaded (accessed 27 April 2022). Measuring the metrics procedure in detail was as follows: The training database includes 999 sentence questions and answers was divided into two datasets(i)_training (699 question and answer pairs) and 300 validation (300 questions and answers pairs). We calculated loss functions. using the Adam optimizer and the loss function binary_crossentropy. After that, we tested the function of the saved model on the test dataset (699 questions and answers) and finally we validated the model performance, using the validation dataset(699 questions and answers)

### 2.3 Case study

The objective of the case study is to repurpose a compound that could target the inflammatory process regulated by Th17 infiltration of the brain during the depression. The constraints on the characteristic of the repurposed drug include the ability to inhibit Th17 differentiation, enhance depression prognosis and pass through the blood-brain barrier as well as being biologically safe ^8,11^. The question used to query the system is “What are anti-oxidant drugs”. The output from our model was further filtered by examining the drug’s oral bioavailability and physicochemical properties. Bioavailability was calculated using the molinspiration online program (www.molinspiration.com). Following Lipinski’s rule for oral absorption of compounds, we calculated the compounds’ molecular weight, number of hydrogen bond donors and acceptors, the topological polar surface area, and number of rotatable bonds, where two violations of this rule will result in the poor oral absorption. clogP, as well as solubility, mol-weight, Tpsa, drug-likeness, and Drug score, was determined by employing Osiris Property Explorer (OPE; http://www.organic-chemistry.org/prog/peo/), The toxicity for each drug was calculated the probability for mutagenicity, tumorigenicity, Irritant-ability and reproductive effectiveness through using the same server. We utilized the BBB predictor (http://www.cbligand.org/BBB) to predict the ability of all the repurposed compounds to cross the blood-brain barrier.

### 2.4 Availability and implementation

To facilitate the use of Adera, we have developed a user python3 application that is quick and reliable. The application with user instruction pages could be found on the Github repository at https://github.com/michel-phylo/ADERA1.

## 3. Results

### 3.1. Workflow

Adera1.1 is a python based neural network workflow specially designed to repurpose drugs pertaining to the field of immunology. Adera1.1 consists of two networks (figure 1). The first network is a universal google encoder designed to convert text into embedded matrices. The second network was designed by us to measure the relevance and sort matching sentences to a given question. The loss of the second network is less than 0.001.

**Figure 1.**
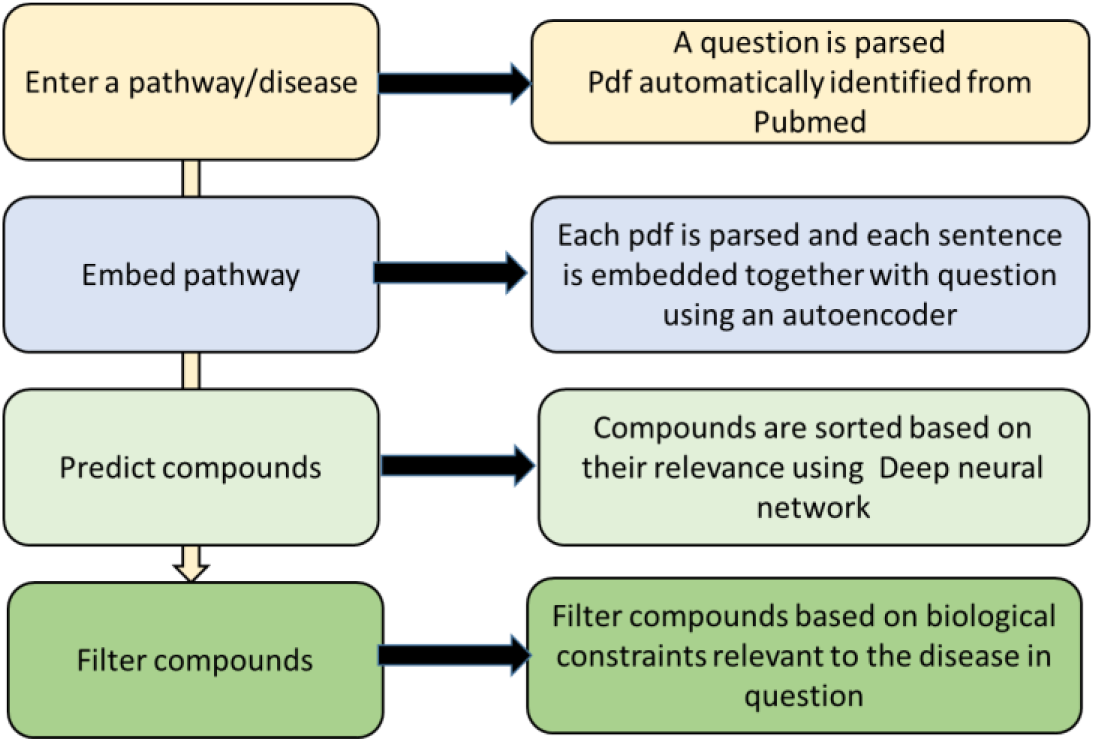
Detailed workflow of our proposal. There are 4 stages in the workflow. The user first chooses or enters a disease, a primary pathway to be investigated. The search step is done to select relevant PubMed articles. Then each accepted article is sorted and parsed. The first network converts each sentence in each PDF to a single embedding matrix. The second neural network sorts each sentence in each pdf based on its relevance to the query. After that, the compounds are filtered based on custom biological constraints.

### 3.2 Accuracy of the first neural network is around 75%

The first neural network is an adaption of the Universal Sentence Encoder ^10^. Using this network each sentence is converted into a matrix of 1*512 (figure 2). The encoder is based on the structure of a deep averaging neural networks. DAN converted words into token embedding before averaging them, then the results is passed through a feedforward deep neural network (DNN) to produce sentence embedding. DAN accuracy is estimated to be around 75%.

**Figure 2.**
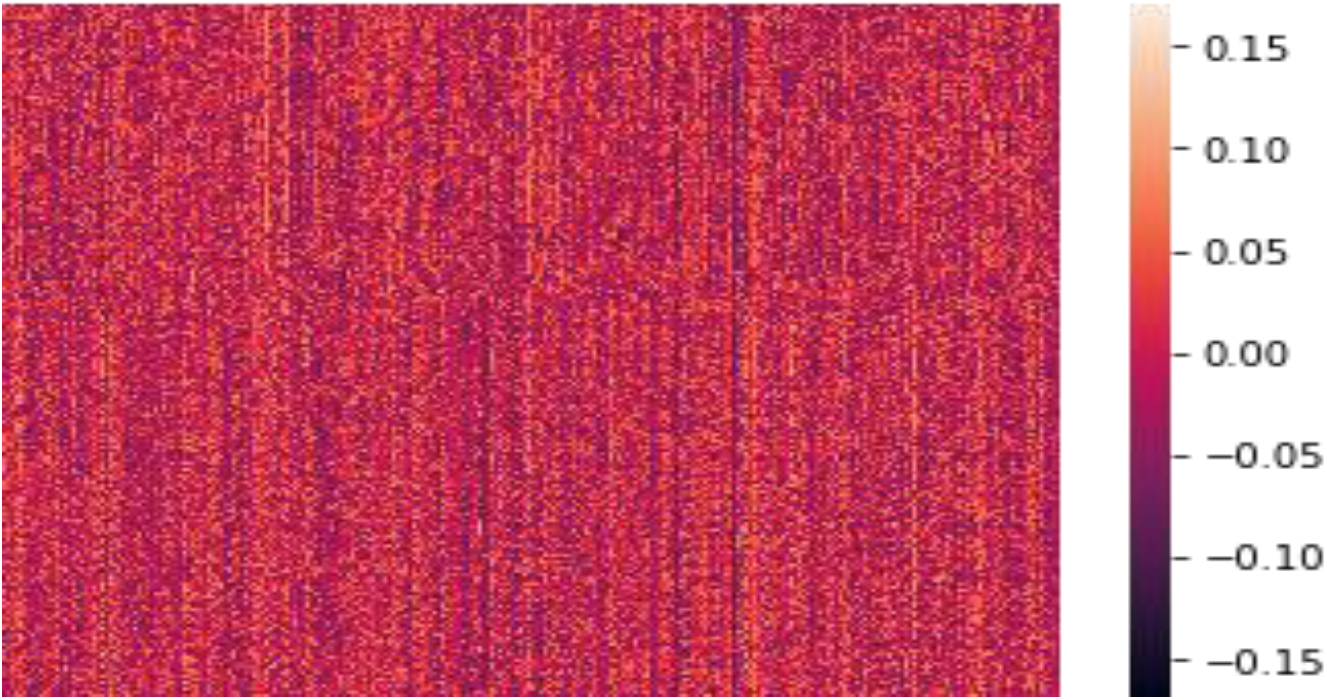
Embedding results of the first network. Using Google universal sentence encoder, we calculated a matrix of (1*512) for each sentence in each pdf including the query question.

**Figure 2.**
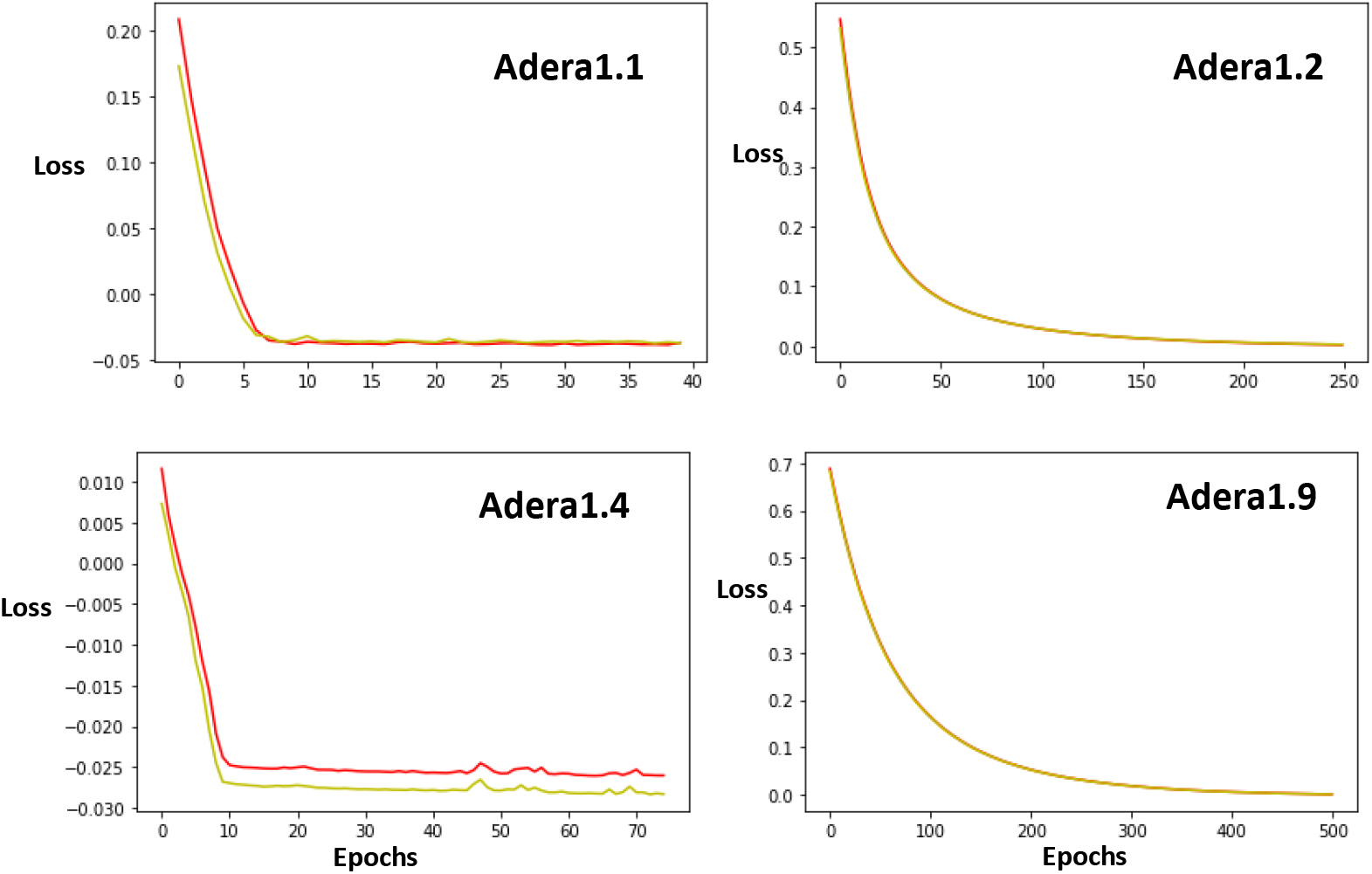
The loss function for the Adera1 network. We explored ten different architectures for relevance calculations. We found that loss using Adera 1.2 outperforms the rest of the networks Green is used for the loss function, while red is for the validation loss calculations.

### 3.3 Output of the second network is dependent on the network architecture

The second neural network’s main function is to sort sentences from each pdf based on their relevance to the question posed (e.g., query). We explored the performance of two networks using 10 different architectures for implementing the relevance network (Table 1). We found that there is a large variation in the network performance based on calculating performance metrics using the SciQ dataset. For the first network Adera1, we calculated the binary cross-entropy loss, while for Adera2, we calculated MSE loss as well as accuracy. In the case of Adera1, we found that the best performing architecture is Adera 1.2, with a loss value of 0.001 (figure 2) (Table 2) and it contains 3 layers; a dense layer with activation of sigmoid, coupled with two successive 2D convolution network, with sigmoid followed by a softmax activation function. Several other structures did not converge such as Adera1.4 which consists of a single dense layer with a relu activation function. Furthermore, the worst-performing network was Adera1.9 with a loss function of 0.01. In the case of Adera2, we found that the best performing is Adera2.4 which is composed of a shallow Relu network (figure 3) (Table3).

**Table 1.**
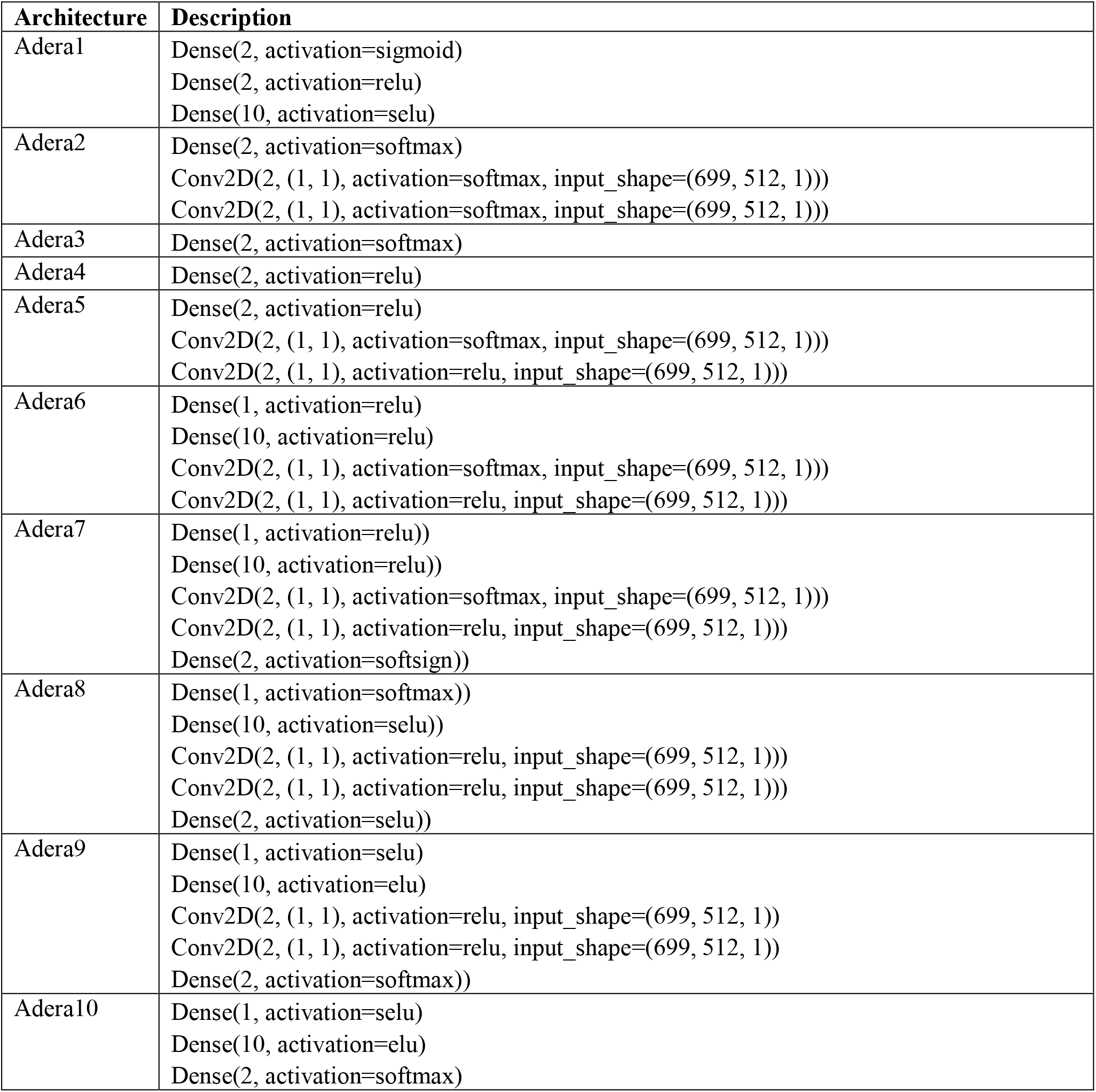
Different structure performance used to calculate relevance.

| **Architecture** | **Description** |
| --- | --- |
| Adera1 | Dense(2, activation=sigmoid)<br>Dense(2, activation=relu)<br>Dense(10, activation=selu) |
| Adera2 | Dense(2, activation=softmax)<br>Conv2D(2, (1, 1), activation=softmax, input_shape=(699, 512, 1)))<br>Conv2D(2, (1, 1), activation=softmax, input_shape=(699, 512, 1))) |
| Adera3 | Dense(2, activation=softmax) |
| Adera4 | Dense(2, activation=relu) |
| Adera5 | Dense(2, activation=relu)<br>Conv2D(2, (1, 1), activation=softmax, input_shape=(699, 512, 1)))<br>Conv2D(2, (1, 1), activation=relu, input_shape=(699, 512, 1))) |
| Adera6 | Dense(1, activation=relu)<br>Dense(10, activation=relu)<br>Conv2D(2, (1, 1), activation=softmax, input_shape=(699, 512, 1)))<br>Conv2D(2, (1, 1), activation=relu, input_shape=(699, 512, 1))) |
| Adera7 | Dense(1, activation=relu))<br>Dense(10, activation=relu))<br>Conv2D(2, (1, 1), activation=softmax, input_shape=(699, 512, 1)))<br>Conv2D(2, (1, 1), activation=relu, input_shape=(699, 512, 1)))<br>Dense(2, activation=softsign)) |
| Adera8 | Dense(1, activation=softmax))<br>Dense(10, activation=selu))<br>Conv2D(2, (1, 1), activation=relu, input_shape=(699, 512, 1)))<br>Conv2D(2, (1, 1), activation=relu, input_shape=(699, 512, 1)))<br>Dense(2, activation=selu)) |
| Adera9 | Dense(1, activation=selu)<br>Dense(10, activation=elu)<br>Conv2D(2, (1, 1), activation=relu, input_shape=(699, 512, 1))<br>Conv2D(2, (1, 1), activation=relu, input_shape=(699, 512, 1))<br>Dense(2, activation=softmax)) |
| Adera10 | Dense(1, activation=selu)<br>Dense(10, activation=elu)<br>Dense(2, activation=softmax) |

**Figure 3.**
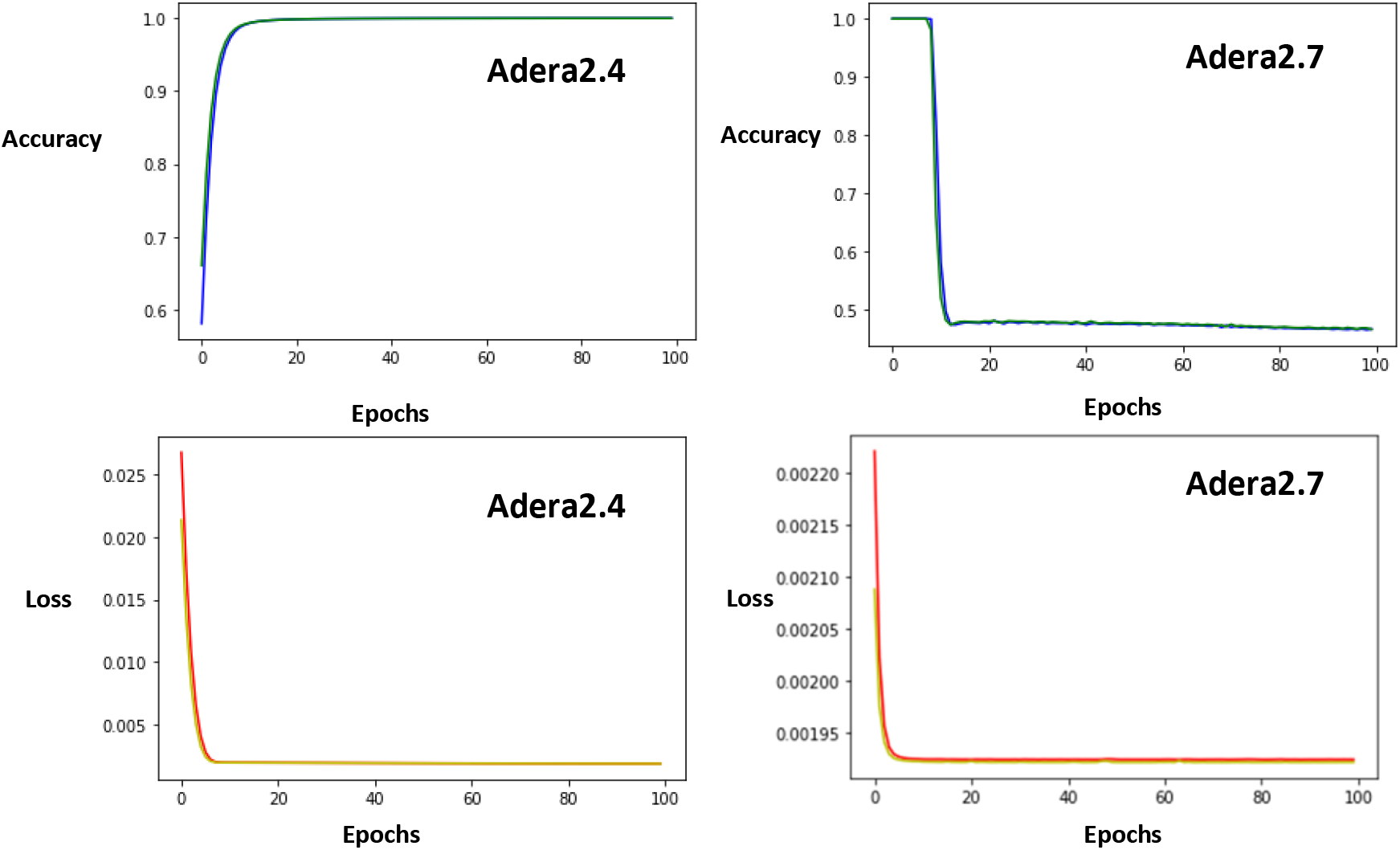
Loss and accuracy function of the Adera2 network. Although Adera2.7 starts with 98% accuracy, it loses this accuracy as the loss function decreases. Conversely, the shallow network of Adera 2.4seems to show both low loss and high accuracy.

**Table 2.** Loss function calculated on using binary cross entropy.

|  | Train dataset |  | Test dataset | Validate |
| --- | --- | --- | --- | --- |
| Architecture | Loss | Val loss | loss | loss |
| Adera1 | -0.0374 | -0.037 | -0.037 | -0.037 |
| Adera2 | -0.001 | -0.001 | -0.001 | -0.001 |
| Adera3 | 0.010 | 0.010 | 0.001 | 0.015 |
| Adera4 | -0.026 | -0.027 | -0.026 | -0.026 |
| Adera5 | -0.020 | -0.02 | -0.02 | -0.02 |
| Adera6 | 0.005 | 0.005 | 0.005 | 0.005 |
| Adera7 | 0.005 | 0.0053 | 0.005 | 0.005 |
| Adera8 | -0.005 | -0.0057 | -0.005 | -0.005 |
| Adera9 | 0.010 | 0.01 | 0.010 | 0.010 |
| Adera10 | -0.006 | -0.006 | -0.006 | -0.006 |

**Table 3.** Loss and accuracy function calculated using MSE.

|  | Train dataset |  |  |  | Test dataset |  | Validate |  |
| --- | --- | --- | --- | --- | --- | --- | --- | --- |
| Architecture | Loss | Accuracy | Val loss | Val acc | loss | accuracy | loss | accuracy |
| Adera1 | 0.001 | 0.361 | 0.001 | 0.301 | 0.0018 | 0.345 | 0.001 | 0.350 |
| Adera2 | 0.252 | 0.960 | 0.252 | 0.960 | 0.252 | 0.505 | 0.250 | 0.505 |
| Adera3 | 0.252 | 0.503 | 0.252 | 0.516 | 0.252 | 0.648 | 0.252 | 0.598 |
| Adera4 | 0.001 | 0.999 | 0.001 | 0.999 | 0.001 | 0.999 | 0.001 | 0.999 |
| Adera5 | 0.002 | 1.000 | 0.002 | 1.000 | 0.002 | 1.000 | 0.002 | 1.000 |
| Adera6 | 0.002 | 0.999 | 0.002 | 0.999 | 0.002 | 0.999 | 0.002 | 0.999 |
| Adera7 | 0.001 | 0.494 | 0.001 | 0.495 | 0.001 | 0.498 | 0.001 | 0.496 |
| Adera8 | 0.002 | 1.000 | 0.002 | 1.000 | 0.002 | 1.000 | 0.002 | 1.000 |
| Adera9 | 0.252 | 0.587 | 0.252 | 0.0001 | 0.252 | 0.549 | 0.252 | 0.503 |
| Adera10 | 0.252 | 0.525 | 0.252 | 0.594 | 0.252 | 0.386 | 0.252 | 0.404 |

### 3.4 *In silico* validation demonstrates Adera performance

We demonstrated the ability of our workflow to repurpose drugs, by its ability to repurpose compounds based on their capabilities to regulate Th17 function in depression. A query question was used to initiate the workflow. The query question was given: “what are anti-oxidant drugs”. The workflow produced ten compounds (Table 4) It could be noticed that the compounds belong to different chemical and physical families. For example, Coumarin is considered a phenylpropanoids^12^ that is used by plants to fend off animals. It is mainly used as an anti-coagulant. On the other hand, Glycyrrhetinic acid is a triterpenoid derived from the root of Liquorice and could function as an anti-cancer and anti-inflammatory drug^13^.

**Table 4.**
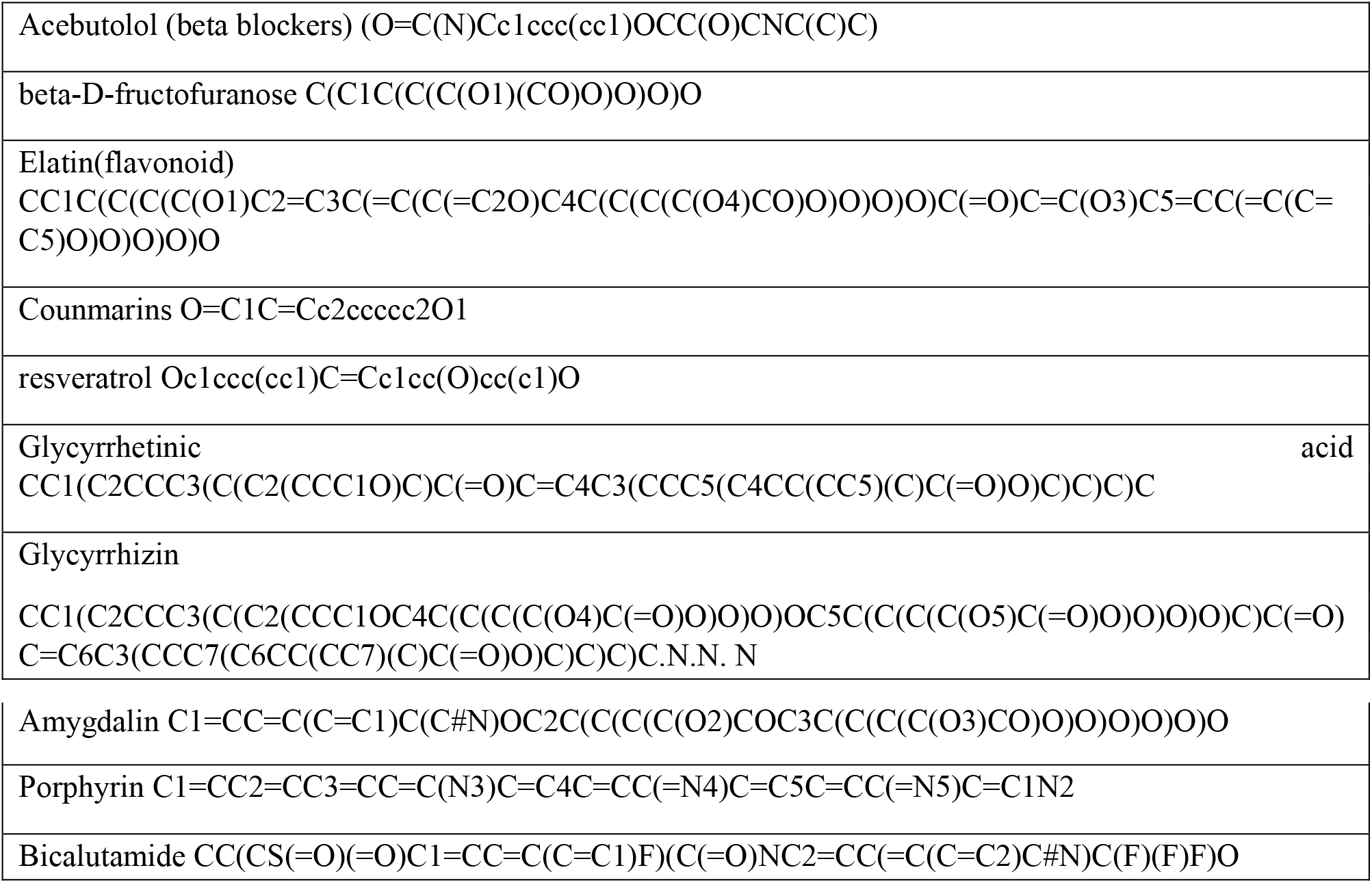
20 compounds were identified using our software.

To examine the feasibility of our predictions to be used as Th17 inhibitors in depression, we subjected the compounds to 11 constraints. In the first set of constraints, pathogenicity was measured (Table 5). Several compounds were predicted not to have any pathogenicity such as Acebutolol, Glyncyrrhetinic acid, glycyrrhizin, Porphyrin, Bicalutamide, and Fucoxanthin. Conversely, Amygdalin is predicted to be of high risk in three of the four categories, highlighting the limitations of our workflow to be inherently aware of toxicity. Moreover, our investigation of the physical and chemical properties of the repurposed compounds revealed that various compounds did not break any of the rules Lipinski’s such as Acebutolol, beta-D-fructofuranose, Coumarin, resveratrol, Bicalutamide, while Glycyrrhizin and Flavonoids broke 3 deeming them weak on oral absorbance. Furthermore, as our repurposing tasks are related to the ability of the compound to inhibit pathogenic Th17 function in the brain during the depression, one of the main characteristics of the required drugs is blood-brain barrier infiltration ability. Our investigation of the Coumarin, resveratrol, Porphyrin, and Bicalutamide is able to cross the BBB (Table 6 and figure 4).

**Table 5.**
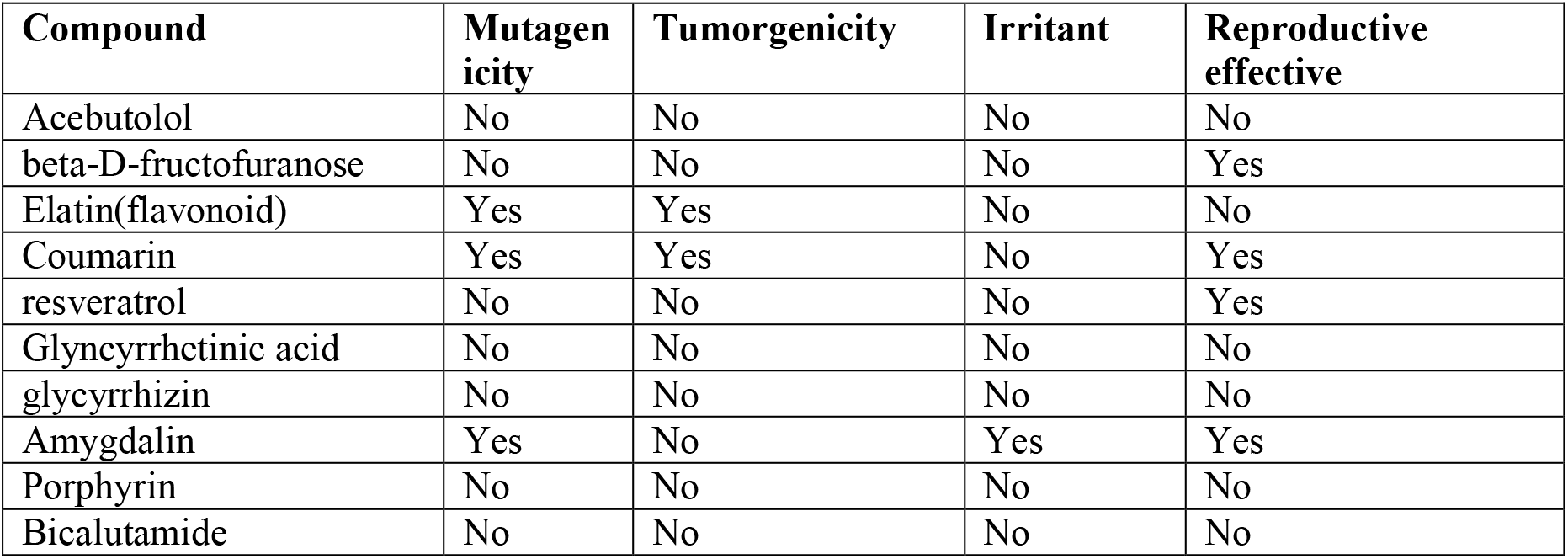
Pathogenicity constraints predication for the identified compounds.

**Table 6.** Comparison of repurposed drugs based on their physical and chemical characteristics.

| Compound | clog | Solubility | Molweight | Tpsa | Druglikness | Drug score | BBB | violations |
| --- | --- | --- | --- | --- | --- | --- | --- | --- |
| Acebutolol | 1.7 | -3.5 | 336.0 | 87.66 | 4.91 | 0.83 | BBB- | 0 |
| beta-D-fructofuranose | -2.7 | 0.38 | 180 | 110 | -2.56 | 0.32 | BBB- | 0 |
| Elatin(flavonoid) | -1.91 | -1.89 | 594 | 267.2 | 0.42 | 0.14 | BBB- | 3 |
| Coumarin | 1.5 | -2.37 | 146 | 26 | -1.83 | 0.12 | BBB+ | 0 |
| resveratrol | 2.83 | -2.86 | 228.0 | 60.69 | -3.25 | 0.27 | BBB+ | 0 |
| Glycyrrhetic acid | 5.36 | -5.78 | 470 | 74.6 | -2.36 | 0.2 | BBB- | 1 |
| Glycyrrhizin | 0.39 | -5.14 | 822 | 267.0 | -4.29 | 0.19 | BBB- | 3 |
| Amygdalin | -3.08 | -1.12 | 457 | 202.3 | -8.7 | 0.09 | BBB- | 2 |
| Porphyrin | 2.05 | -4.34 | 310.0 | 52.54 | 0.97 | 0.67 | BBB+ | 1 |
| Bicalutamide | 2.14 | -5.08 | 430 | 115 | -11 | 0.24 | BBB+ | 0 |

**Table 7.**
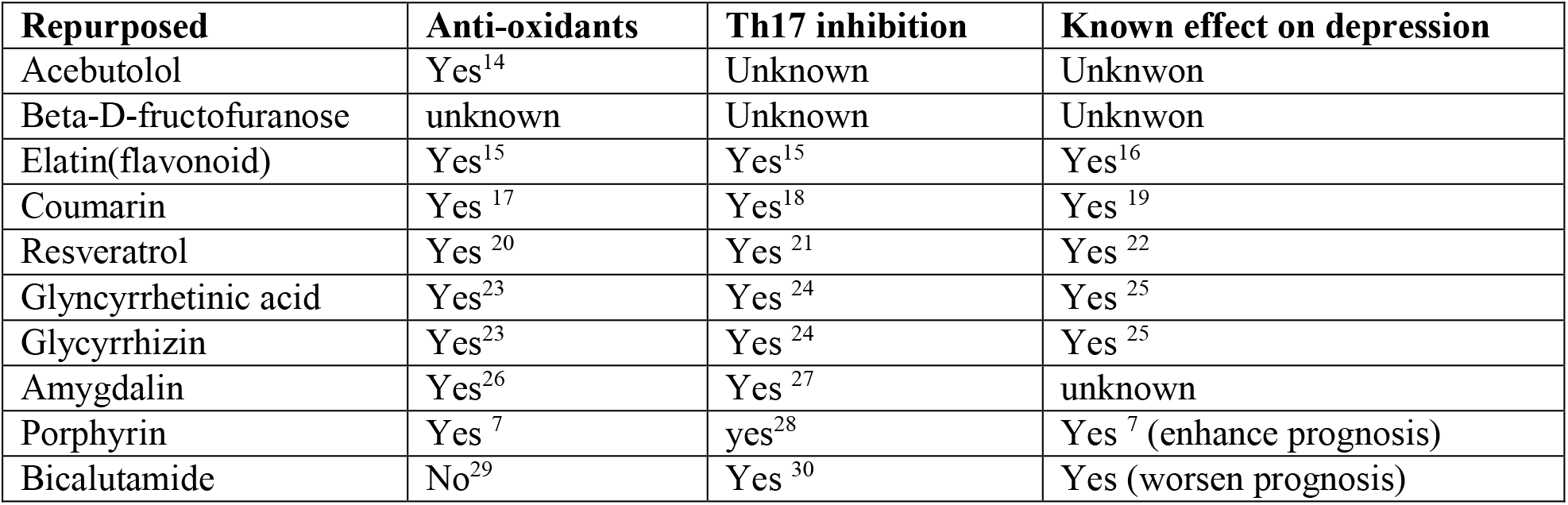

**Figure 4.**
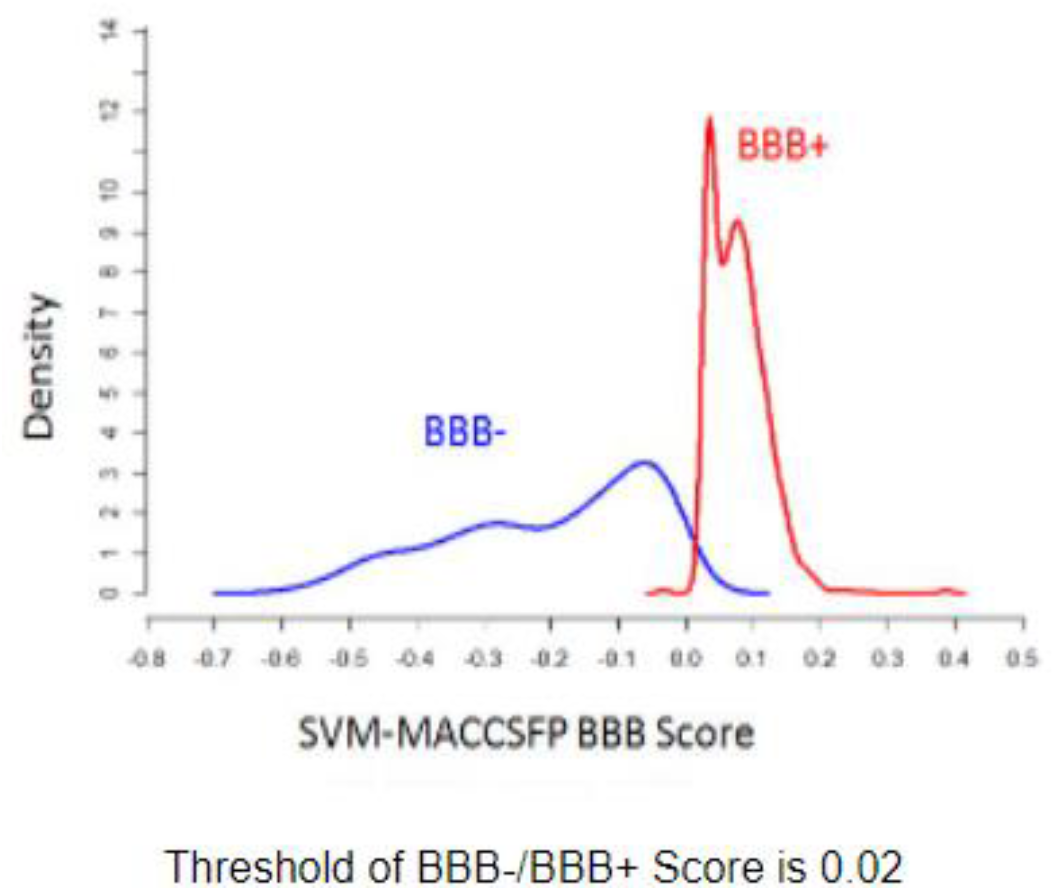
BBB infiltration predictions of Bicalutamide. Bicalutamide scored 0.02 on the SVM-MACCCSFP BBB score indicating that it is highly likely to be able to cross the BBB.

## Discussion

Our system offers a drug repurposing workflow based on the relevance. Thousands of literature is being added to PubMed and similar repositories each year. Mining drug databases is a viable option for drug repurposing. However this approach lacks awareness of the context between the drug and its vast applications and effects that are unlikely to be reported in a single database. Additionally, each day tens of hundreds of novel findings are being reported, thus the feasibility of having access to all of them in a single drug database is low. Additionally our workflow bypass the traditional limitations of NLP approaches by using two deep neural networks that are to reduce false positives.

## Output of the second network highlight importance of architecture

Our approach as well as others demonstrate neural network prediction results are highly dependable on the architecture used as well as the loss function. In this report, we explored the outcome of two-loss functions (i) Binary cross-entropy and (ii) MSE. In the case of the binary loss function, Adera 2 given by a Dense layer with an activation input of a softmax, followed by two 2D convolution layers with an activation input of softmax performs best. Whereas, the highest loss was associated with using a dense layer with a relu activation function followed by a layer of 2d convolution layer with a softmax and a 2d convolution layer with a relu activation function. Thus it seems that using the softmax activation function with binary cross-entropy might be yielding the lowest loss. The reason behind this phenomenon could be based on the observation that, binary cross-entropy is a form of measuring loss-based KL-divergence (relative entropy)^31^. KL divergence computes the difference between two probability distributions (computed and expected). Relu (rectified linear unit) output is equal to the input if the input is positive otherwise it is zero^32^. Although Relu avoids vanishing gradients, however, it can produce zero value gradients when the activation is less than zero, thus preventing the weights from being adjusted^33^. This leads to having constant KL-divergence values (e.g., no convergence). On the other hand, softmax is better at handling multiple classes and is often used as the last layer. Softmax computes the exponential of each element of the input and normalizes it by dividing its value by the summation of the exponentials of all the inputs^34^. The main advantage of the softmax is that when the input is a negative value, the function output is never negative. This ensures that each class will be associated with a single probability and that the sum of probabilities will be one. This in turn ensures that KL divergence values are updated leading to better loss values. Notably, in the case of the MSE function, Adera4 scored the highest accuracy (0.98%) and a reasonable loss (0.018). Adera4 is a shallow network consisting of a single dense layer with a Relu activation function. Importantly it performed much better than a single dense layer with softmax (Adera3). The reason behind that is the MSE function given by the square value of the error ensures the predicted weight is never zero in the case of Relu^35,36^.

## Case study results shows our workflow predictions accuracy

We demonstrated the ability of our workflow to correctly repurpose a compound that can target Th17 expansion in the brain. We have previously shown that Th17 is capable of infiltrating the brain during depression d causing a localized inflammation of the hypothalamus^37^. This inflammation is associated with depression-like behavior^38^. Currently, depression drugs focus on the neurotransmitter aspect of the disease and overlook the inflammatory side. Thus repurposing drugs to target the inflammatory symptoms of depression can help ease the symptoms and ensures a better quality of life. We used Adera to mine Pubmed Pdf for anti-oxidant drugs. We investigated a list of ten drugs for their ability to fulfill the needed objective (i.e., target Th17 expansion in the brain during the depression). Additionally, we subjected the drugs to a rigorous list of constraints that ensures the repurposed drug would fulfill the objective while being biologically safe, orally available, and nontoxic. Our investigation proposes the use of Porphyrin and Bicalutamide. Our findings are supported by the literature where it was shown that both drugs have an inhibitory effect on Th17 and enhance depression prognosis^7,28,30^.

## Limitations and future direction

At the moment Adera 1.1 is unaware of the protein structure of the repurposed drug’s target. Text mining of NGS public repositories including RNA-seq (bulk and single cell) as well as Chip-seq as well as forming text miners that would be able to predict drug toxicity and interactions are highly achievable. Automatic filtering of repurposed drugs based on their characteristics could prove efficient in reducing R&D costs^39^.

## Conclusion

Our workflow Adera 1.1 is capable of text mine highly specific Pdfs to search for a specific question. Adera1.1 high accuracy of 98% is based on its ability to predict the relevance of each sentence in the pdf compared with posed query using a relevance dedicated neural network. Adera usage could reduce the time and the cost needed for R&D of drugs discovery.

## Acknowledgement

We would like to Thanks Macrious Abraham and Meriam Joachim for their fruitful discussions and inspirations. We would like to thank the team of Nlet and COST for their financial support.

## References

1. Scannell, J. W., Blanckley, A., Boldon, H. & Warrington, B. Diagnosing the decline in pharmaceutical R&amp;D efficiency. Nat. Rev. Drug Discov. 11, 191–200 (2012).

2. Prasad, V. & Mailankody, S. Research and development spending to bring a single cancer drug to market and revenues after approval. JAMA Intern. Med. (2017) doi:10.1001/jamainternmed.2017.3601.

3. Kubick, N., Pajares, M., Enache, I., Manda, G. & Mickael, M.-E. Repurposing Zileuton as a Depression Drug Using an AI and In Vitro Approach. Molecules 25, 2155 (2020).

4. Anstey, A. & Lear, J. T. Azathioprine: Clinical pharmacology and current indications in autoimmune disorders. BioDrugs (1998) doi:10.2165/00063030-199809010-00004.

5. Nag, S. et al. Deep learning tools for advancing drug discovery and development. 3 Biotech 12, 110 (123AD).

6. Lotfi Shahreza, M., Ghadiri, N., Mousavi, S. R., Varshosaz, J. & Green, J. R. A review of network-based approaches to drug repositioning. Brief. Bioinform. 19, 878–892 (2018).

7. Corsello, S. M. et al. The Drug Repurposing Hub: A next-generation drug library and information resource. Nature Medicine (2017) doi:10.1038/nm.4306.

8. Mickael, M. E. et al. RORγt-Expressing Pathogenic CD4 + T Cells Cause Brain Inflammation during Chronic Colitis. J. Immunol. 208, 2054–2066 (2022).

9. Mickael, M.-E. et al. Paracellular and Transcellular Leukocytes Diapedesis Are Divergent but Interconnected Evolutionary Events. Genes (Basel). 12, 254 (2021).

10. Cer, D. et al. Universal Sentence Encoder. https://tfhub.dev/google/.

11. Bhaumik, S. & Basu, R. Cellular and molecular dynamics of Th17 differentiation and its developmental plasticity in the intestinal immune response. Frontiers in Immunology (2017) doi:10.3389/fimmu.2017.00254.

12. Jacobowitz, J. R. & Weng, J. K. Exploring Uncharted Territories of Plant Specialized Metabolism in the Postgenomic Era. Annual Review of Plant Biology (2020) doi: 10.1146/annurev-arplant-081519-035634.

13. Graebin, C. S. The Pharmacological Activities of Glycyrrhizinic Acid (“Glycyrrhizin”) and Glycyrrhetinic Acid. in Reference Series in Phytochemistry (2018). doi: 10.1007/978-3-319-27027-2_15.

14. Krouf, D. et al. Changes in serum lipids and antioxidant status in west Algerian patients with essential hypertension treated with acebutolol compared to healthy subjects. Med. Sci. Monit. (2003).

15. Al-Khayri, J. M. et al. Flavonoids as Potential Anti-Inflammatory Molecules: A Review. Molecules 27, 2901 (2022).

16. Pannu, A., Sharma, P. C., Thakur, V. K. & Goyal, R. K. Emerging role of flavonoids as the treatment of depression. Biomolecules (2021) doi: 10.3390/biom11121825.

17. Kostova, I. et al. Coumarins as Antioxidants. Curr. Med. Chem. (2012) doi:10.2174/092986711803414395.

18. Yao, R. et al. Regulatory effect of daphnetin, a coumarin extracted from Daphne odora, on the balance of Treg and Th17 in collagen-induced arthritis. Eur. J. Pharmacol. (2011) doi:10.1016/j.ejphar.2011.08.019.

19. Capra, J. C. et al. Antidepressant-like effect of scopoletin, a coumarin isolated from Polygala sabulosa (Polygalaceae) in mice: Evidence for the involvement of monoaminergic systems. Eur. J. Pharmacol. (2010) doi:10.1016/j.ejphar.2010.06.043.

20. Xia, N., Daiber, A., Förstermann, U. & Li, H. Antioxidant effects of resveratrol in the cardiovascular system. British Journal of Pharmacology (2017) doi:10.1111/bph.13492.

21. Guo, N. H. et al. The potential therapeutic benefit of resveratrol on Th17/Treg imbalance in immune thrombocytopenic purpura. Int. Immunopharmacol. (2019) doi:10.1016/j.intimp.2019.04.061.

22. Moore, A., Beidler, J. & Hong, M. Y. Resveratrol and depression in animal models: A systematic review of the biological mechanisms. Molecules (2018) doi:10.3390/molecules23092197.

23. Li, X. L., Zhou, A. G., Zhang, L. & Chen, W. J. Antioxidant status and immune activity of glycyrrhizin in allergic rhinitis mice. Int. J. Mol. Sci. (2011) doi: 10.3390/ijms12020905.

24. Chen, X. et al. Glycyrrhizin ameliorates experimental colitis through attenuating interleukin-17-producing T cell responses via regulating antigen-presenting cells. Immunol. Res. (2017) doi:10.1007/s12026-017-8894-2.

25. Murck, H. et al. Adjunct Therapy With Glycyrrhiza Glabra Rapidly Improves Outcome in Depression—A Pilot Study to Support 11-Beta-Hydroxysteroid Dehydrogenase Type 2 Inhibition as a New Target. Front. Psychiatry (2020) doi:10.3389/fpsyt.2020.605949.

26. Albogami, S. et al. Evaluation of the effective dose of amygdalin for the improvement of antioxidant gene expression and suppression of oxidative damage in mice. PeerJ (2020) doi:10.7717/peerj.9232.

27. Gago-López, N., Lagunas Arnal, C., Perez, J. J. & Wagner, E. F. Topical application of an amygdalin analogue reduces inflammation and keratinocyte proliferation in a psoriasis mouse model. Exp. Dermatol. (2021) doi: 10.1111/exd.14390.

28. Zhang, Y., Zhang, L., Wu, J., Di, C. & Xia, Z. Heme Oxygenase-1 Exerts a Protective Role in Ovalbumin-induced Neutrophilic Airway Inflammation by Inhibiting Th17 Cell-mediated Immune Response. J. Biol. Chem. 288, 34612–34626 (2013).

29. Chen, K. C. et al. Bicalutamide elicits renal damage by causing mitochondrial dysfunction via ROS damage and upregulation of HIF-1α. Int. J. Mol. Sci. (2020) doi:10.3390/ijms21093400.

30. Zhong, S., Huang, C., Chen, Z., Chen, Z. & Luo, J. L. Targeting inflammatory signaling in prostate cancer castration resistance. J. Clin. Med. (2021) doi:10.3390/jcm10215000.

31. Bu, Y., Zou, S., Liang, Y. & Veeravalli, V. V. Estimation of KL Divergence: Optimal Minimax Rate. in IEEE Transactions on Information Theory (2018). doi:10.1109/TIT.2018.2805844.

32. Zeiler, M. D. et al. On rectified linear units for speech processing. in ICASSP, IEEE International Conference on Acoustics, Speech and Signal Processing-Proceedings (2013). doi:10.1109/ICASSP.2013.6638312.

33. Liu, M., Chen, L., Du, X., Jin, L. & Shang, M. Activated Gradients for Deep Neural Networks. IEEE Trans. Neural Networks Learn. Syst. (2021) doi:10.1109/tnnls.2021.3106044.

34. Liang, X., Wang, X., Lei, Z., Liao, S. & Li, S. Z. Soft-Margin Softmax for Deep Classification. in Lecture Notes in Computer Science (including subseries Lecture Notes in Artificial Intelligence and Lecture Notes in Bioinformatics) (2017). doi:10.1007/978-3-319-70096-0_43.

35. Mickael, M. E., Heydari, A., Crouch, R. & Johnstone, S. Estimation of stress-strain relationships in vascular walls using multi-layer hyperelastic modelling approach. in Computing in Cardiology vol. 37 (2010).

36. Mickael, M. Modelling Baroreceptors Function. (2012).

37. Mickael, M. E. et al. RORγt-Expressing Pathogenic CD4+T Cells Cause Brain Inflammation During Chronic Colitis. bioRxiv 2021.09.01.458634 (2021) doi:10.1101/2021.09.01.458634.

38. Beurel, E. & Lowell, J. A. Th17 cells in depression. Brain. Behav. Immun. 69, 28–34 (2018).

39. Kubick, N., Henckell Flournoy, P. C., Klimovich, P., Manda, G. & Mickael, M. E. What has single-cell RNA sequencing revealed about microglial neuroimmunology? Immunity, Inflammation and Disease (2020) doi: 10.1002/iid3.362.

